# Information- and Communication-Centric Approach in Cell Metabolism for Analyzing Behavior of Microbial Communities

**DOI:** 10.1101/2023.08.23.554558

**Authors:** Zahmeeth Sakkaff, Andrew Freiburger, Nidhi Gupta, Massimiliano Pierobon, Christopher S. Henry

**Affiliations:** Argonne National Laboratory, Lemont, IL, USA; Department of Computer Science and Engineering, University of Nebraska, Lincoln, NE, USA; University of Chicago, Chicago, IL, USA

## Abstract

Microorganisms naturally form community ecosystems to improve fitness in diverse environments and conduct otherwise intractable processes. Microbial communities are therefore central to biogeochemical cycling, human health, agricultural productivity, and technologies as nuanced as nanotechnology-enabled devices; however, the combinatorial scaling of exchanges with the environment that predicate community functions are experimentally untenable. Several computational tools have been presented to capture these exchanges, yet, no attempt has been made to understand the total information flow to a community from its environment. We therefore adapted a recently developed model for singular organisms, which blends molecular communication and the Shannon Information theory to quantify information flow, to communities and exemplify this expanded model on idealized communities: one of *Escherichia coli* (*E. coli*) and *Pseudomonas fluorescens* to emulate an ecological community and the other of *Bacteroides thetaiotaomicron* (*B. theta*) and *Kleb Ciella* to emulate a human microbiome interaction. Each of these sample communities exhibit critical syntrophy in certain environmental conditions, which should be evident through our community mutual information model. We further explored alternative frameworks for constructing community genome-scale metabolic models (GEMs) – mixed-bag and compartmentalized. Our study revealed that information flow is greater through communities than isolated models, and that the mixed-bag framework conducts greater information flow than the compartmentalized framework for community GEMs, presumably because the latter is encumbered with transport reactions that are absent in the former. This community Mutual Information model is furthermore wrapped as a KBase Application (<u>*R*</u>*un <u>F</u>lux <u>M</u>utual Information Analysis, RFMIA*) for optimum accessibility to biological investigators. We anticipate that this unique quantitative approach to consider information flow through metabolic systems will accelerate both basic and applied discovery in diverse biological fields.

**Author Summary:** Microorganisms frequently communicate information via information-bearing molecules, which must be fundamentally understood to engineer biological cells that properly engage with their environments, such as the envisioned Internet of Bio-NanoThings. The study of these molecular communications has employed information and communication theory to analyze the exchanged information via chemical reactions and molecular transport. We introduce an information- and communication-centric computational approach to estimate the information flow in biological cells and its impacts on the behavior of single organisms and communities. This study complements our previous work of cell metabolism by developing an end-to-end perspective of molecular communication based on enzyme-regulated reactions. We explore the mutual information using Shannon information theory, measured in bits, between influential nutrients and cellular growth rate. The developed *RFMIA* computational tool is deployed in the U.S. Department of Energy’s Systems Biology Knowledgebase, where it quantitatively estimates information flow in both organism and community metabolic networks and extends recent developments in computer communications to explore and explain a new biology for the open-source community.

## Introduction

Microorganisms naturally assemble into communities to diversify strengths and weakness and to conduct complex functions such as biogeochemical cycling, intricate bioproduction, and digesting intractable nutrients. Cellular metabolism is the foundation of these transformative processes, where chemical reactions convert environmental substrates into energy and biomass in a tightly regulated pipeline [1] (Fig. 1). These reactions are activated by the cell’s environment to optimally survive, often meaning growth and reproduction [2]. Environmental perturbations may therefore be useful to control cellular behaviors, although the limits of this approach and the communication mechanisms by which cells and communities coordinate functions in response to environmental triggers remain opaque. The specific mediating molecules (e.g. quorum sensing) and triggers for these communication mechanisms represent a large knowledge gap [3, 4] around cellular communication. Community communications especially remain understudied [5], partly due to the difficulty in recovering representative samples [6]; hence computational biology is an attractive alternative domain to study communities without necessitating a complete experimental description of a system. Some computational models apply top-down ecological principles to represent microbial communities while other models represent communities from the bottom-up assembly of metabolic pathways and fundamental biochemistry [7]. Molecular communication (MC) in Fig. 1 is an ideal blend of both approaches – while also leveraging mathematics, computer science, and chemical engineering – to fully capture the multi-dimensional complexity of community interactions by deducing information flow from chemical exchanges. We previously derived an MC abstraction of a single communication channel of interdependent inputs and outputs [8], where information propagates from media into cellular metabolism and where we derived quantitative limits for this information flow [9], and for a two molecular communication channels for each community [10]: one where regulatory mechanisms depend upon combinations of extracellular compounds, and the other where cell metabolism depends on growth and extracellular exchange. The application of information theory in these papers (illustrated in Fig. 2) expresses the performance of both channels and examined the end-to-end (E2E) limits to molecular communication system, such as exchange flux and biomass growth.

**Fig 1.**
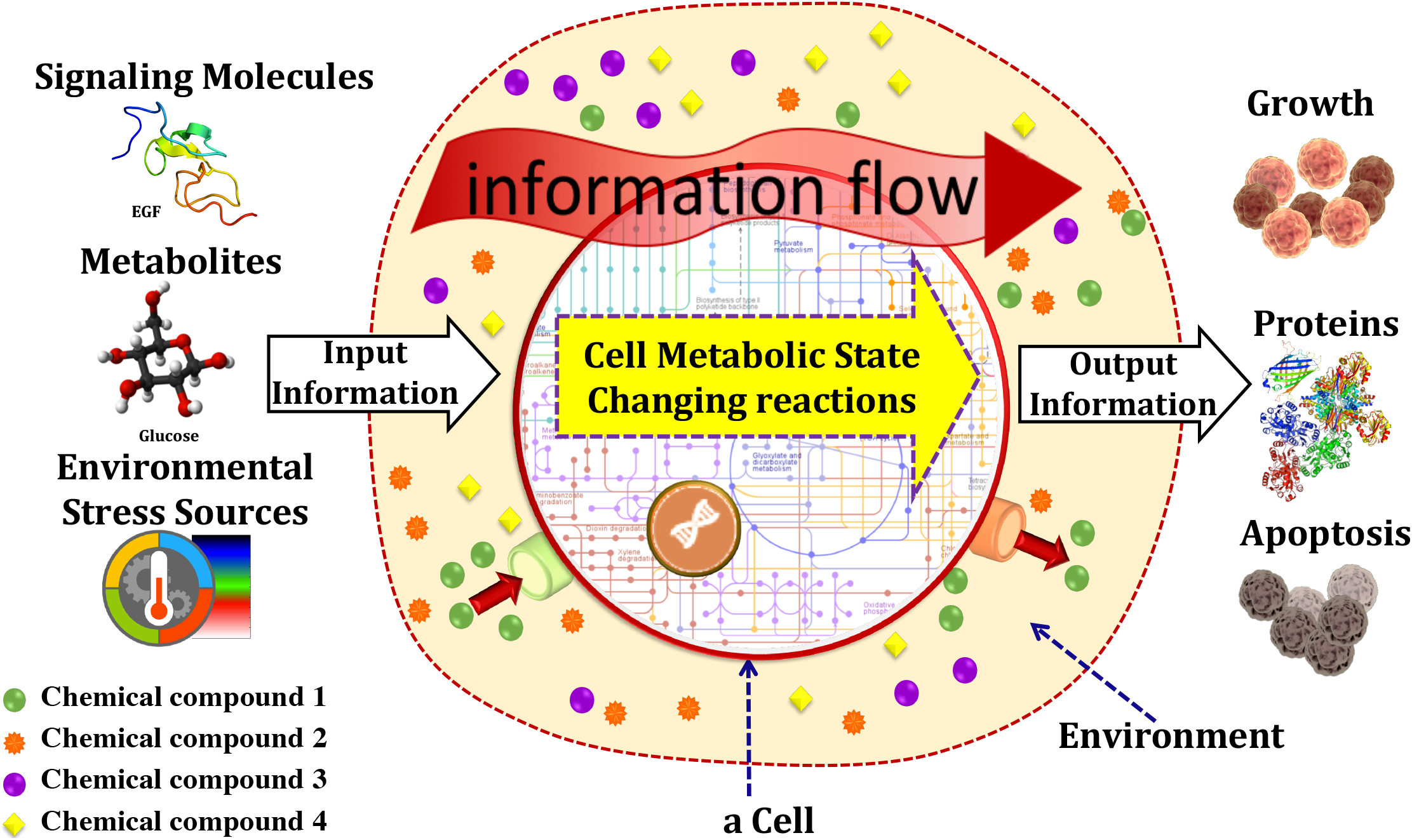
Molecular communication systems send and receive information through chemical reactions. Information flows from the cell’s environment in the form of input signals such as signaling molecules, metabolites, environmental stress sources like temperature, and pH. Output information flows from the metabolic state and how information of the internal cell state can be perceived from the outside environment in the form of growth of a cell, proteins and apoptosis of a cell.

**Fig 2.**
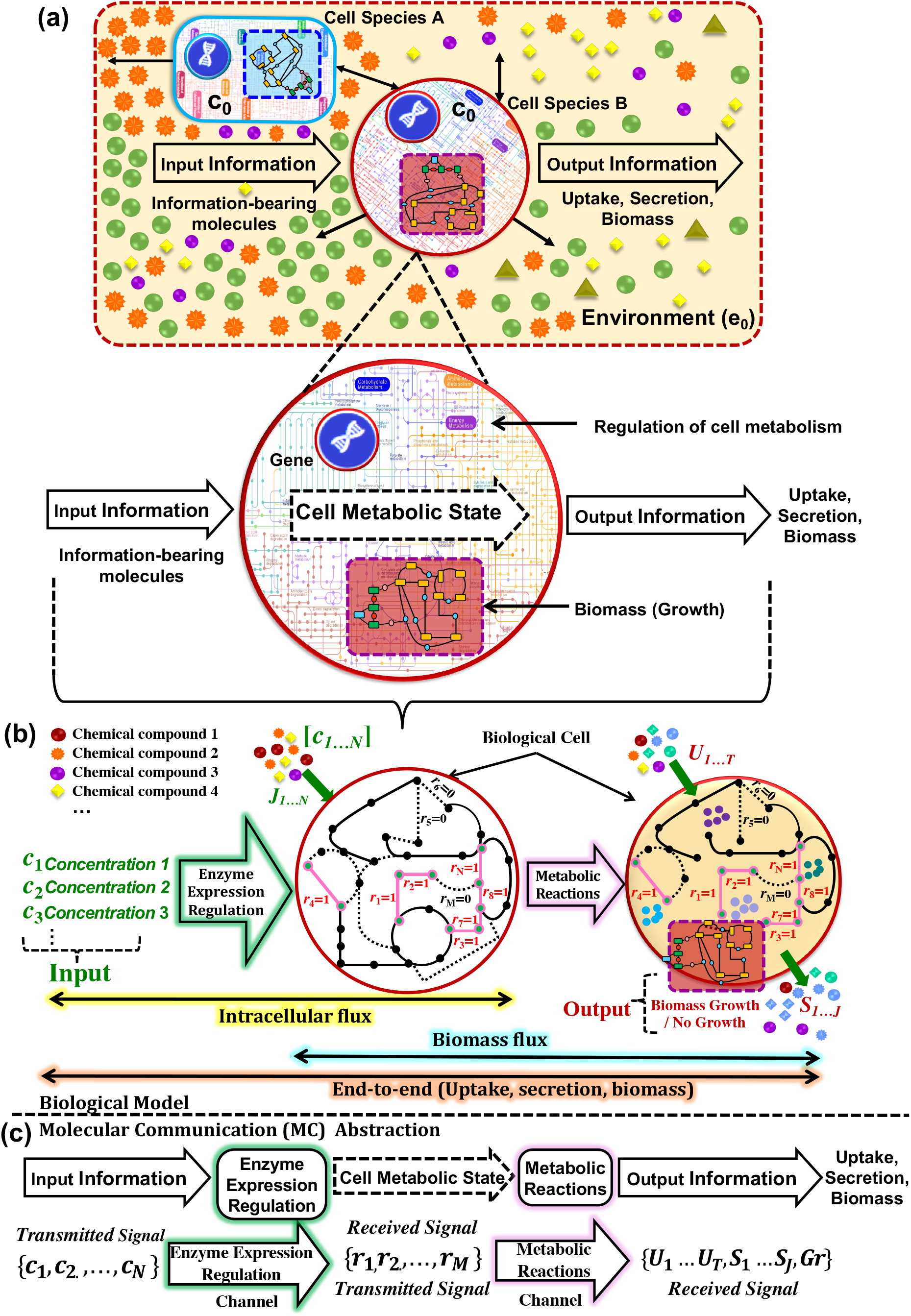
Overall graphical representation of the molecular communication abstraction of cell metabolism. Interaction of two different cell species A, and B with the presence of various compounds in the environment and the flow of information. (b) The cell takes certain concentrations of chemical compounds into the cell metabolism, where the variations in these concentrations cause state changes in the cell’s metabolic network. Intracellular flux represents the cells response to the chemical composition of the cell’s environment; Biomass flux captures variations in the growth rate of the cell in response to environmental changes; and End-to-End describes variations in the uptake and secretion of chemical compounds and the biomass (cell growth), which can be measured when we consider the cell metabolic processes from the perspective of the environment. (c) Mapping MC abstraction with biological model.

Herein, we expand our previously defined molecular communication model into a Mutual Information score (MI) that is crafted to examine community systems. with different combinations of seven substrates for two human gut microbes – an obligate anaerobe bacterium *B. theta* and an Archaea *M. smithii*, where the substrates were found to propagate different amounts of information [**?**]. The MI score uniquely 1) characterizes cellular and inter-species molecule exchanges, 2) detects extracellular signatures – exchange fluxes and biomass growth – that indicate intra-cellular information flow, and 3) generates data that can reveal community interactions and thereby minimize resource-intensive experiments to elucidate community communications and their sensitivity to environmental perturbations. MI in Fig. 2 leverages Shannon information theory [11, 12] to quantify variability in metabolic information flow and steady-state FBA [13] to define an upper limit of information flow from the given environment based on metabolic flux, biomass growth, and E2E (uptake, secretion of metabolites and biomass). Shannon’s theory defines information as the details that distinguish a state of a system from the space of possible system states, which is depicted in Fig. 3. The MI score is the uncertainty difference between the uncertainty contained in the input molecule(s) and the conditional uncertainty of the channel output, and thereby quantifies information change via each channel. We exemplify the MI score to understand an idealized microbiome community of *Escherichia coli* (*E. coli*) and *Bacteroides thetaiotaomicron* (*B. theta*) that has a prominent role on our adaptive immunity [14] and biotic health [15], and which exhibits a unique community synergy of *B. theta* only growing in aerobic environments when coupled with *E. coli* [16]. We moreover compared the information differences between two alternative frameworks for defining community GEMs: 1) the mixed-bag framework, where all member networks are pooled into a single network; and (2) the compartmentalized framework, where each member network is segregated into a designated compartment that can exchange with a common environment compartment. We consistently observe a higher MI score in communities, especially the mixed-bag abstraction, relative to isolates, which is qualitatively expected yet quantitatively invaluable when comparing different community designs or ecological factors that govern community formation. The MI score therefore has far-reaching implications in diverse biological fields and should accelerate engineering efforts of controlling microbiomes or synthesizing communities. The MI score is available as a point-and-click Application (*Run Flux Mutual Information Analysis, RFMIA* in the U.S. Department of Energy’s Systems Biology Knowledgebase (KBase) [17] where genomes can be constructed into metabolic networks and ultimately merged community models [18] (See the example KBase Narrative: https://kbase.us/n/40576/330/).

**Fig 3.**
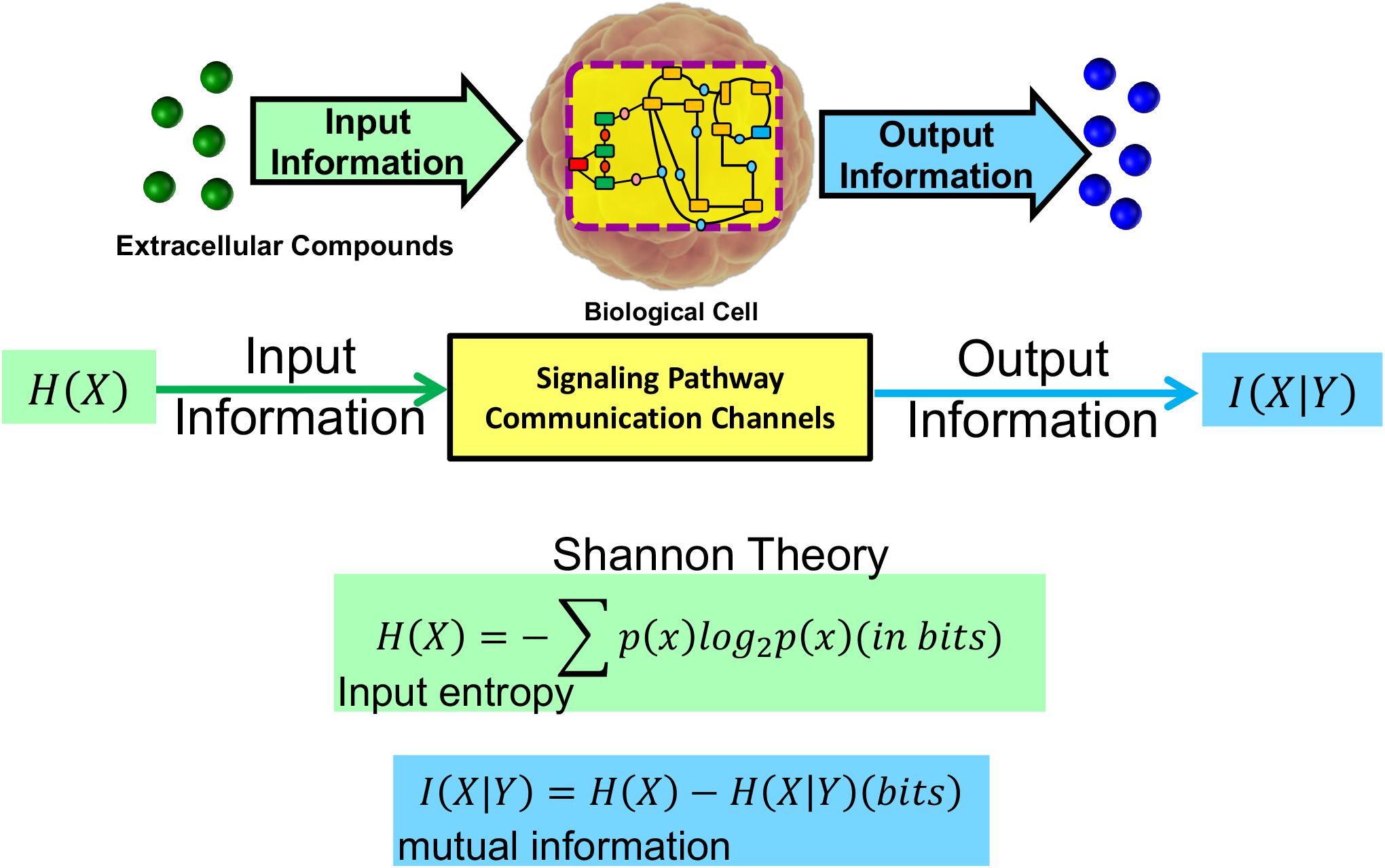
Fundamental mathematical framework of Shannon information theory in the biological system. Formulation of pure uncertainty of the input signals presented as extracellular compounds and the mutual information for output signals presented as cellular responses to inputs.

## Materials and Methods

Mutual Information (*I*) is defined as the input uncertainty minus the output uncertainty that depends on the inputs: *I* = *H*(*inputs*) − *H*(*inputs, outputs*). The MI was applied to explore how much environmental perturbations (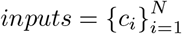, as the concentration of all *N* examined substrates) can control metabolic flux and biomass growth 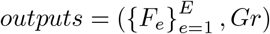, as all exchange fluxes and the biomass growth flux [10], respectively). The MI for the overall E2E channel quantifies information flow that can be perceived from the environment through exchange fluxes and biomass growth

### Upper Bound to the Steady-State Mutual Information – UBS-sMI

FBA is employed to computationally evaluate (1) and (3), which yields the profile of activated reactions in a given the environment and enables estimating the state 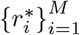 that maximizes biomass production and the associated fluxes *i*.*e*., 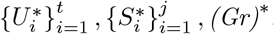. Parsimonious FBA (pFBA) [19] is furthermore applied to reduce the space of optimal solutions to those with the lowest total flux, thereby imposing a secondary optimization of metabolic efficiency while improving reproducibility of the predicted flux profiles. The upper bound for *MI* of metabolic flux [9]:

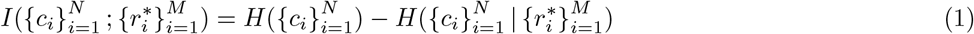

where {*c*_*i*_}_*i*=1_ is the initial concentrations. The relations between *MI* and its upper bound

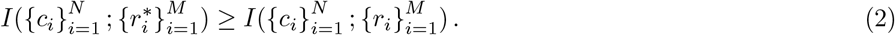

and *MI* of biomass growth and its upper bound

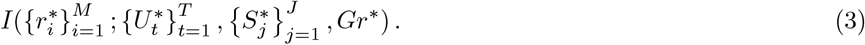

are analogously derived. The upper bound of the E2E steady-state mutual information 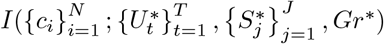 is solved by substituting 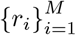 with 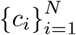 and then integrating. The mathematical formulation is presented in supplementary **S1**.

### Reconstruction of Community Models

We sourced published GEMs for *E. coli*/**iML1515** [20] and *B. theta*/**iAH991** [21] that offer the most complete and experimentally-validated reconstructions to-date for these organisms. These member GEMs were assembled to a mixed-bag community model – the pooled amalgamation of all reactions – and as a compartmentalized community model – each GEM is assigned a unique compartment and all compartments exchange with a common extracellular compartment – which are contrasted in Fig. 4. These community model frameworks are functionally identical in FBA, with the difference that transport reactions are not present in mixed-bag community models since they lack compartments. The **Merge metabolic models into community model** tool in KBase constructed each version of community model for the two-member community.

**Fig 4.**
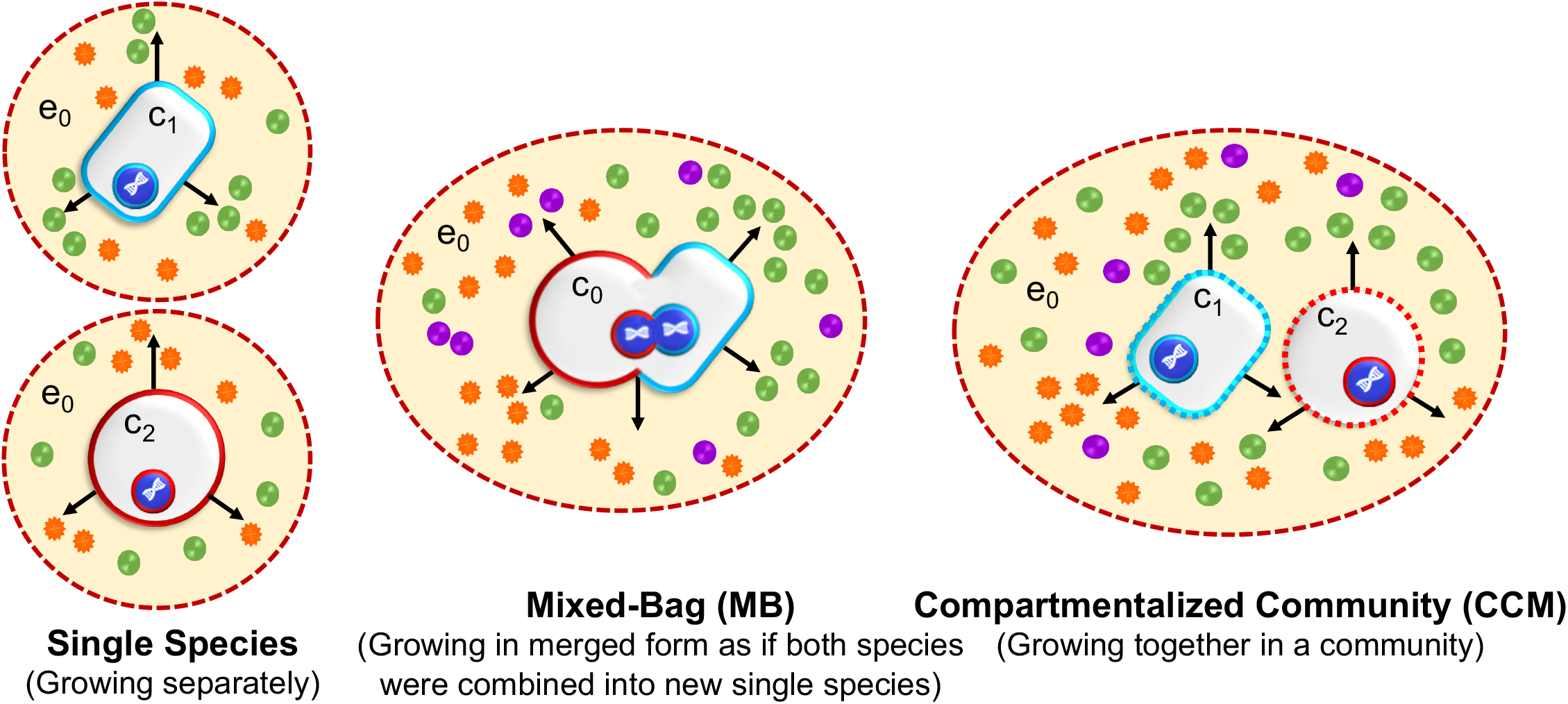
Details of interspecies models in this study. Three different formalisms we use to study information flow through metabolic systems. Left-hand models show the environmental interaction of individual species. Middle MB shows individual species blended into a single new species. Right hand CCM, where individual species tightly interact with each other while interacting with the environment. Species 1 are represented in purple, species 2 in green, and anything common in both organisms in orange. Filled and open circles represent extracellular and intracellular metabolites, respectively; solid boxes mean that compartments outside of which the quasi-steady state assumption may not hold, and dashed boxes indicate compartments outside of which the steady-state assumption continues to hold. As stated, the communities did not have equal abundances, and that situation may lead to inaccurate conclusions about any interactions. Hence, the relative abundance values allow one to formulate the biomass composition for the entire community.

### Mutual Information score

The MI is a metric in units of bits for the metabolic ability to process information from a given environment, where the activity of state-changing chemical reactions are either activated (1) or inactive (0) depending on the media. The MI score is proportional with information flow between the cell metabolism and its environment.

## Results

### Integration of the Mutual Information Approach in KBase

The *RFMIA* KBase Application in Fig. 6 executes the MI score, where the user-friendly workflow in Fig. 5 accepts inputs of the models, media components, a *mmol* of carbon consumption limit, and model objectives and returns tables and charts that succinctly communicate information flows as well as rank compounds based on their influence on information flow. Steps 1–5 in Fig. 5a,b explain the backend processes of the RFMIA KBase App.

**Fig 5.**
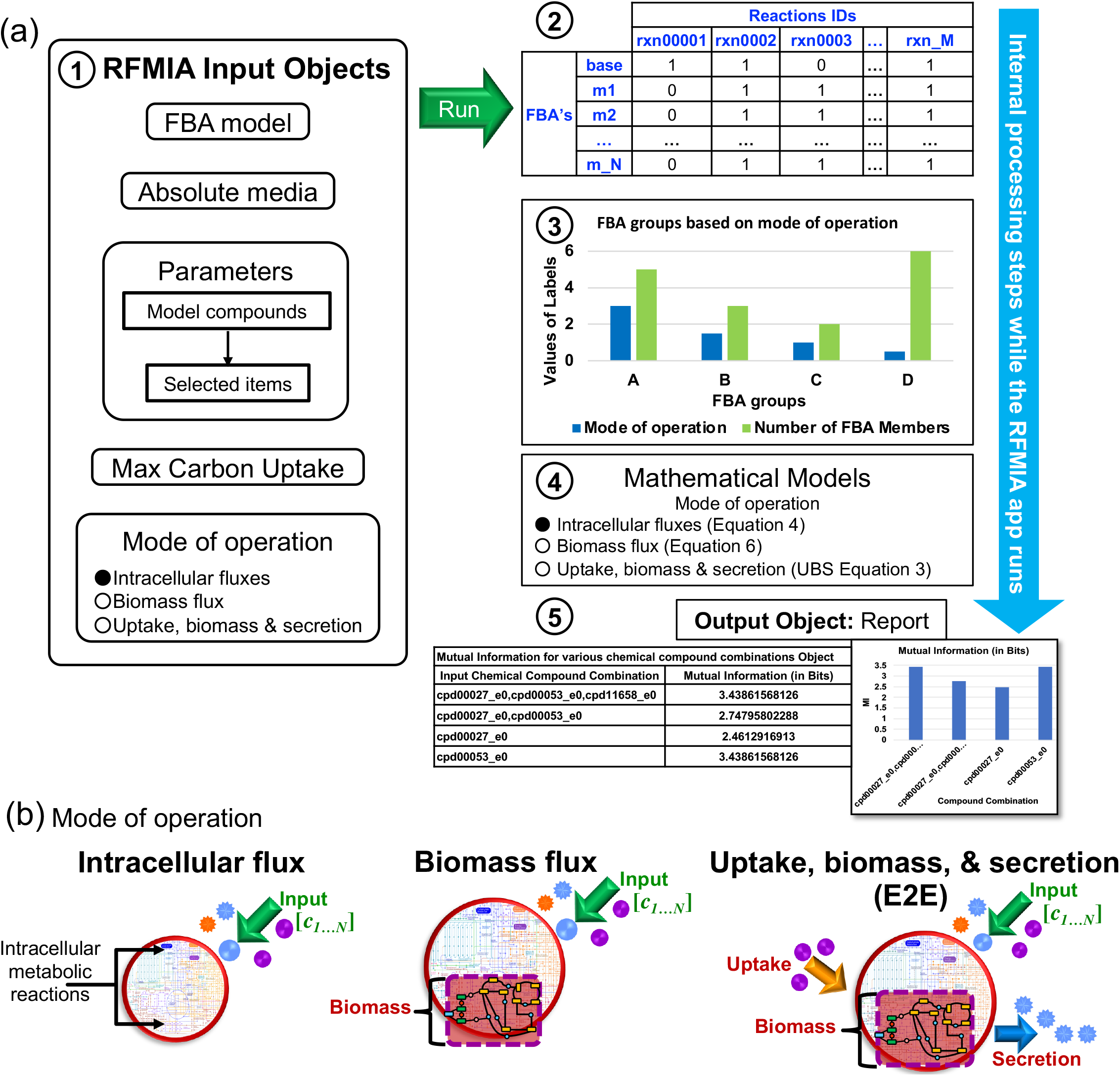
General RFMIA tool pipeline. (a) 1-5 steps involved: 1-input to the tool, 2-internal processing of reaction table with FBAs, 3-FBA groups with MI values, 4-MI math models and calculations, and 5-MI table and bar chart. (b) A pictorial description of the mode of operation.

**Fig 6.**
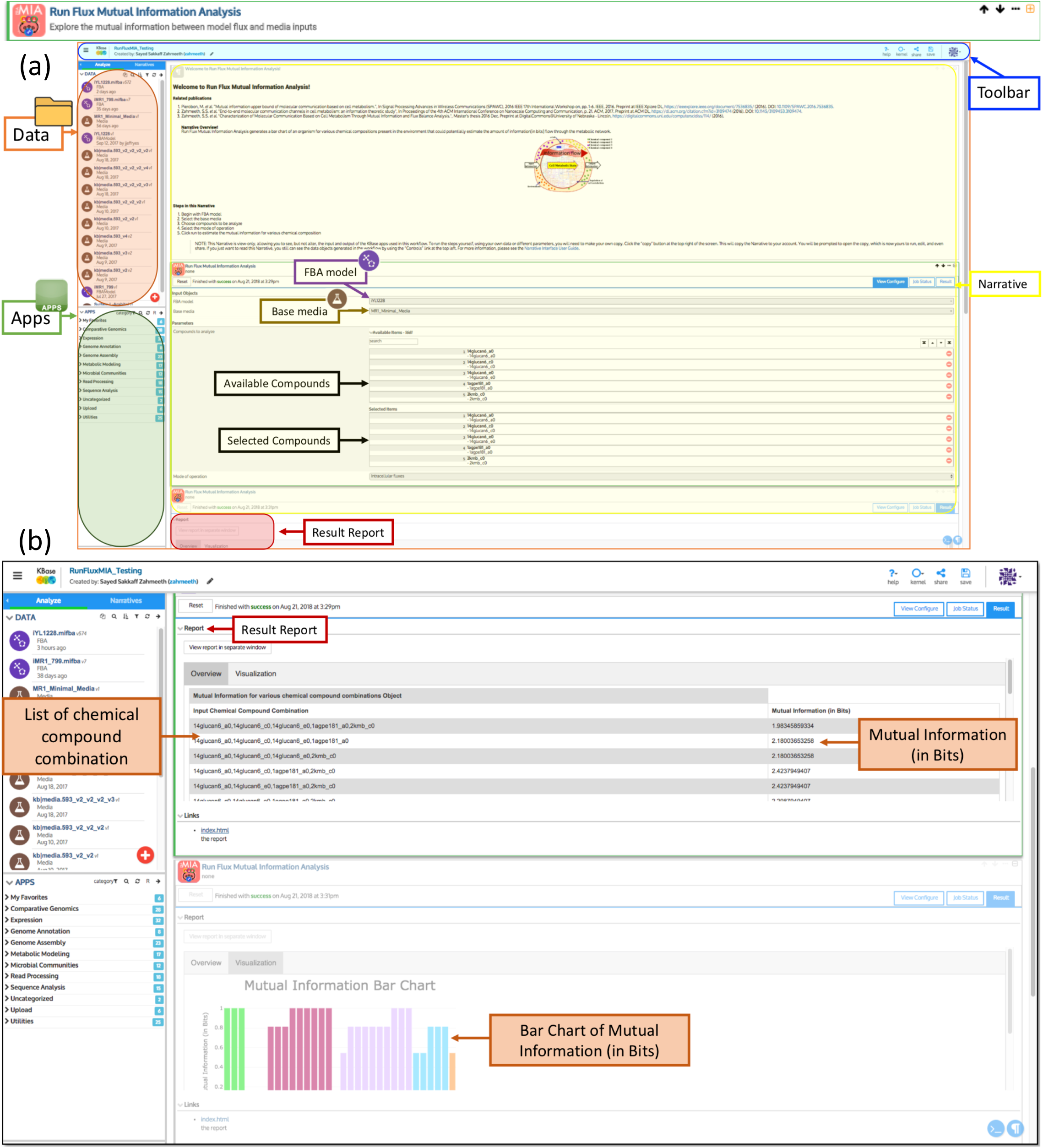
RFMIA online user interface available in KBase [?] as point and click apps. (a) Describes the user interface containing narrative description, RFMIA tool features, input data, and the list of applications. (b) RFMIA run results in tabular and bar chart formats.

### Integration of Published *E. coli* and *B. theta* Models and Combination into Community models

The FBA simulation results, GEM metadata, and growth rates for aerobic and anaerobic conditions among each of the seven input nutrients are tabulated in Table 1 (corollary content is available in **S3** and Narrative https://kbase.us/n/81575/184/). The published individual models were subtly curated with missing pathways that are necessary for our simulation: *E. coli* was curated to produce dextrin from starch and sucrose and *B. theta*/**iAH991V2** was curated by adding choline exchange and transporter reactions and removing sink-chols c0 and sink-hpyr c0 sink reactions (the final models are available in Narrative https://kbase.us/n/40576/330/). We constrain the total number of carbon atoms that a model can consume 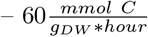 by default, which corresponds to 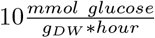 when glucose is the only carbon source – to limit simulated growth to a reasonable value even when several carbon sources are simultaneously utilized. The ecological role of *B. theta* as an aerotolerant – can survive but not grow in aerobic environments [22] – digester of polysaccharides [23] was captured by manually setting *MI* and biomass growth in its GEM to zero when the isolated model was simulated in an aerobic environment, while leaving the GEM unconstrained in a community.

**Table 1.** Individual and community models statistics.

| Model Statistics | Single |  | Community |  |
| --- | --- | --- | --- | --- |
|  | *B. theta* / iAH991 | *E.coli* / iML1515 | MB | CCM |
| Original Model Description |  |  |  |  |
| Number reactions | 1246 | 2379 | 3166 | 3625 |
| Number compounds | 1179 | 1878 | 2398 | 2944 |
| Genes | 993 | 1515 | 2508 | 2508 |
| Number compartments | 2 | 3 | 3 | 5 |
|  | c0-Cytosol<br>e0-Extracellular | c0-Cytosol<br>e0-Extracellular<br>p0-Periplasm | c0-Cytosol<br>e0-Extracellular<br>p1-Periplasm | c0-Cytosol<br>c1-*E. coli*<br>c2-*B. theta*<br>e0-Extracellular<br>p1-Periplasm |
| Aerobic Glucose MM | Glucose carbon source (presence of Oxygen) |  |  |  |
| Active reactions | 0 | 417 | 578 | 1116 |
| Objective value / Growth | 0 | 0.477218 | 0.428449 | 0.414962 |
| Anaerobic Glucose MM | Glucose carbon source (absence of Oxygen) |  |  |  |
| Active reactions | 594 | 413 | 564 | 979 |
| Objective value / Growth | 0.162608 | 0.16799 | 0.428449 | 0.173746 |
| Aerobic Dextrin MM | Glucose replace with Dextrin carbon source (presence of Oxygen) |  |  |  |
| Active reactions | 0 | 0 | 573 | 1042 |
| Objective value / Growth | 0 | 0 | 0.428449 | 0.428449 |
| Anaerobic Dextrin MM | Glucose replace with Dextrin carbon source (absence of Oxygen) |  |  |  |
| Active reactions | 522 | 0 | 569 | 934 |
| Objective value / Growth | 0.214225 | 0 | 0.428449 | 0.428449 |
| Aerobic Glutamine MM | Ammonia replace with Glutamine nitrogen source (presence of Oxygen) |  |  |  |
| Active reactions | 0 | 416 | 563 | 1034 |
| Objective value / Growth | 0 | 1.11357 | 0.428449 | 0.428449 |
| Anaerobic Glutamine MM | Ammonia replace with Glutamine nitrogen source (absence of Oxygen) |  |  |  |
| Active reactions | 537 | 413 | 566 | 982 |
| Objective value / Growth | 0.214225 | 0.675723 | 0.428449 | 0.428449 |
| Aerobic Sulfate MM | Hydrogen Sulfate removed (presence of Oxygen) |  |  |  |
| Active reactions | 0 | 417 | 580 | 1148 |
| Objective value / Growth | 0 | 0.477218 | 0.428449 | 0.411395 |
| Anaerobic Sulfate MM | Hydrogen Sulfate removed (absence of Oxygen) |  |  |  |
| Active reactions | 0 | 413 | 576 | 987 |
| Objective value / Growth | 0 | 0.16799 | 0.428449 | 0.172423 |
| CMM – Compartmentalized Community Model MB – Mixed-Bag MM – Minimal Media (Details in supplementary S3) |  |  |  |  |

Several noteworthy observations are available from the various models in the various media. First, the community models, especially the mixed-bag model, generally exhibited higher growth relative to the individual models, although the highest for growth was observed with *E. coli* in aerobic conditions. Second, the community models successfully grew on dextrin despite that *E. coli* failed to grow as an isolate, which may be explained by *E. coli*’s inability metabolize complex polysaccharides [24] but this is not completely understood [25]. Third, the community models grew on glutamine (a nitrogen source) equivalently to glucose and dextrin (carbon sources), however, *E. coli* nearly doubled and tripled the community growth rate in anaerobic and aerobic conditions on glutamine relative to the carbon sources, respectively. Finally, the community models grew equivalently on sulfate as glucose (the mixed-bag model grew slightly more, while *B. theta* failed to grow and *E. coli* grew slightly more than the community models. The first two observations demonstrate community synergy that may justify its stable formation among these members. The latter two observations reveal conditions where *E. coli* self-sacrifices to join the community, since it could grow more as an independent isolate, notwithstanding that sulfur metabolism in bacteria is poorly understood [26].

### Mutual Information Analysis of Isolate and Community models

The RFMIA App was applied to each model isolate and community model, with upper bounds for all combinations of the seven nutrients in E2E: (1) Glucose (*C*_6_*H*_12_*O*_6_), (2) Glutamine (*C*_5_*H*_10_*N*_2_*O*_3_), (3) Dextrin (*H*(*C*_6_*H*_10_*O*_5_)*xOH*), (4) Ammonia (*NH*_3_), (5) Sulfate 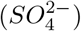, (6) Hydrogen sulfate (*H*_2_*SO*_4_), and (7) Oxygen (*O*_2_). Diverse responses were observed in Fig. 7 for aerobic and anaerobic conditions and between the isolates and community models. Compound combinations without information flow were removed.

**Fig 7.**
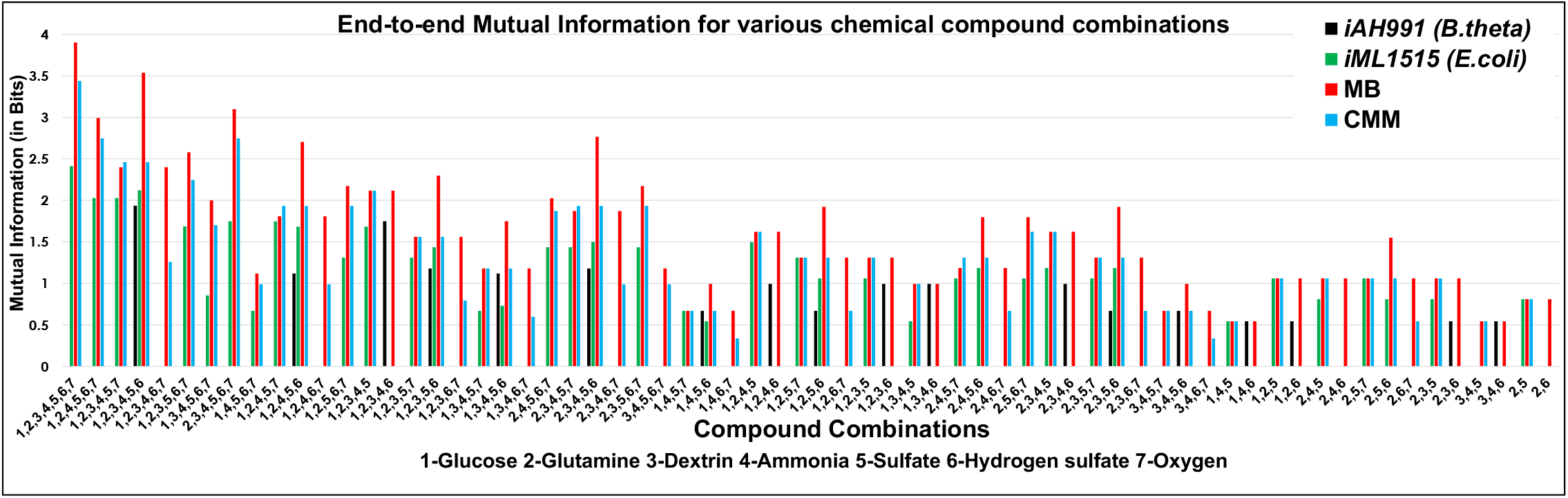
End-to-end (E2E) mutual information chart with respect to various input compound combinations. E2E mutual information for the different combinations of seven compounds of individual *E. coli, B. theta*, mixed-bag(MB), and compartmentalized community model (CCM), with respect to flux of uptaken and secreted compounds and biomass. Compound combinations with zero bits of all four models are not presented. Corresponding bar chart values are available in the supporting document S4.

Figure 7 reveals that both *E. coli* and *B. theta* consist of information flow between 0.544–2.122 bits and 0.669–1.936 bits, where {1,2,3,4,5,6} generates the maximum values for each member with containing two carbon sources, two nitrogen sources, and two sulfur sources and {1,4,5,6} generates the minimum values for each member with one carbon source, one nitrogen source, and two sulfur sources. *B. theta* exhibits greater information flow in anaerobic conditions with the presence of 1-2 sources of carbon, nitrogen, and sulfur: e.g. {1,2,3,4,6}, {1,2,4,6}, {2,3,4,6}, {3,4,5,6}, {1,4,6}, {2,3,6}, and {3,4,6}. *E. coli* generally exhibits greater information flow otherwise, particularly with glutamine, which is consistent with the biological role of the latter as an obligate anaerobe that has substantial limitations as an isolate. The presence of sulfate in combination with carbon and nitrogen sources augments information flow in *E. coli*, while the absence of a sulfur, carbon, or nitrogen source often results in zero information flow.

Almost every set of E2E inputs in Fig. 7 manifests in greater information flow in the community models compared with single species, which supports the ecological synergy of these members. Information flow is even greater in mixed-bag models than compartmentalized models, presumably because the latter is encumbered with transport exchanges between compartments. This trend was observed in both aerobic conditions in Figure 8 and anaerobic conditions in Fig. 9. *E. coli* further outperformed *B. theta* in all conditions except two anaerobic conditions {1,3,4,5,6} and {1,4,5,6}, which is consistent with the ecological roles of each member [14]. Information flow in the absence of a carbon source is zero for *B. theta*, low for *E. coli*, and highest for the community models, which further reveals circumstances that justify community formation. The presence of glucose and dextrin manifested in the greatest information flow, followed by glucose exceeding dextrin when deployed singularly, ceteris paribus. The presence of both glutamine and ammonia nitrogen sources similarly manifested in greater information flow than either source individually, followed by glutamine exceeding ammonia. *B. theta* does not exchange information in the absence of hydrogen sulfate, regardless of the presence of sulfate, which suggests that *B. theta* may require additional protons for survival, while both isolates exhibit zero information flow when hydrogen sulfate is present without sulfate. These results affirm the nutritional necessity of these mineral and energy sources, and indicate that information flow generally follows *B. theta < E. coli <* CCM *<* MB.

**Fig 8.**
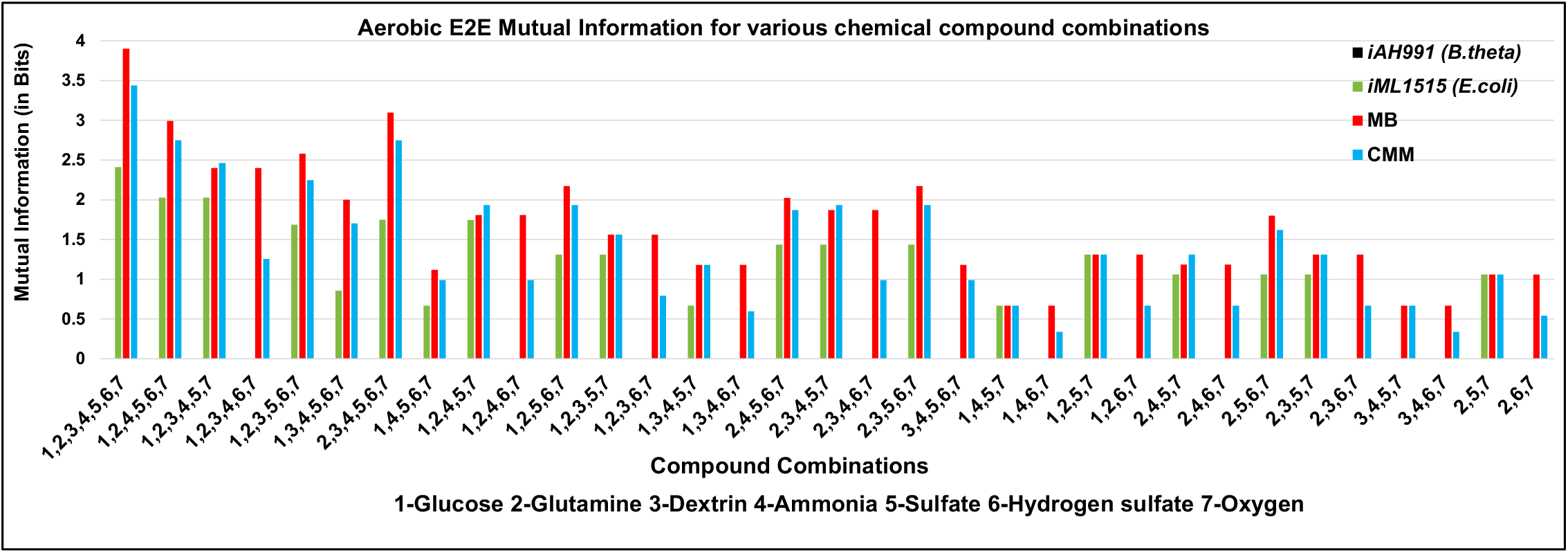
Aerobic E2E mutual information chart for different combinations of seven compounds. MI values of *E. coli, B. theta*/**iAH991V2**, mixed-bag (MB), and compartmentalized community model (CCM) with all the combinations of seven compounds, with respect to E2E MC systems. MI combinations with zero bits are not presented.

**Fig 9.**
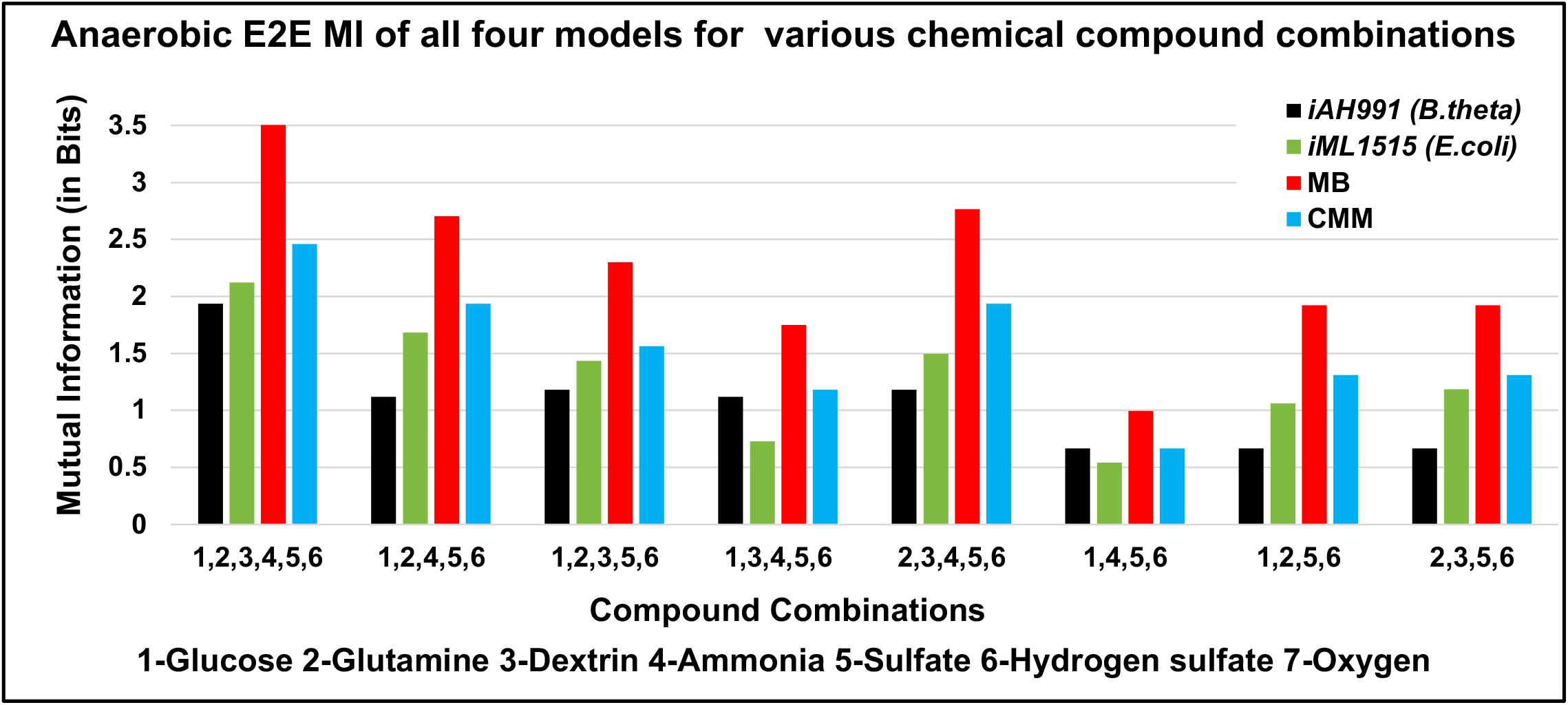
Anaerobic E2E mutual information values of all four models for seven compound combinations. A chart describes the anaerobic E2E MI values of *B. theta*/**iAH991V2**, *E. coli*, mixed-bag (MB), and compartmentalized community model (CCM) with all the combinations of seven compounds respectively, to E2E MC systems. MI combinations with zero bits are not presented.

## Discussion

Biological cells leverage communication, both among themselves and with other cells, to conduct relatively complex cellular functions and govern interactions and community dynamics. Understanding these understudied communication mechanisms is therefore essential for understanding human health, ecological systems, and agricultural productivity, and ultimately for rational engineering a range of cellular behaviors or stable communities. We devised a Mutual Information score (MI) to quantify the information flow of a metabolic system and probe these communication mechanisms via a unique molecular communication model that applies Shannon information theory. We exemplify MI with an idealized 2-member microbiome community of *E. coli* and *B. theta*, where information flow was determined to be greater in community models than in isolated members, consistent with the Shannon Information theory’s definition of information which scales proportionally with the number of possible system states. The mixed-bag community framework furthermore exhibited greater information flow than compartmentalized community models since this framework is not encumbered by transporters to exchange compounds between compartments and may therefore represent a theoretical limit of a frictionless community metabolism. The MI values of *E. coli* were also generally greater than those of *B. theta* – except for several anaerobic conditions – which is consistent with the broader ecological habitable zone of *E. coli*. We further ingrained MI into KBase as an Application (*Run Flux Mutual Information Analysis*) that conveniently visualizes information flows for concise interpretation and importantly streamlines usage of MI and broadens its accessibility to non-technical biologists. MI analyses identify the combinations of nutrients that optimally trigger information flow, and may therefore be a valuable tool to reduce the number of resource-intensive experiments to probe a given metabolic system. We envision that MI and the *RFMIA* App will accelerate basic and applied discoveries in myriad biological fields. Future work will characterize larger microbial communities and chart information flow as a function of member abundances to understand how competitive forces influence information flow in community systems.

## Supporting information

**S1 Mathematical Models. Mutual Information Mathematical Models** Details of the mathematical models are in the Mutual Information MathematicalModels.pdf file.

**S2 Figures 7, 8, and 9**.

**S3 Table Minimal Media (MM)** The table contains the compounds the minimal necessities for growth of the both organisms. Details of the compound list are in file Btheta Ecoli minimal media.csv

**S4 Figures 7, 8, and 9 bar chart values in excel file** The values and the original formats for B theta E Coli BarCharts. are in the Excel file sheets 1, 2, and 3, respectively.

## Acknowledgments

We thank the KBase developers team in Chris Henry’s lab at Argonne National Laboratory (ANL) for their active support throughout the development of this research. Also, special thanks to Dr. Gail W. Pieper at ANL for helpful discussions and comments on revisions. This work was financially supported by the Molecular and Biochemical Telecommunications (MBiTe) Lab at the Department of Computer Science and Engineering, University of Nebraska -Lincoln. This work was supported by the US National Science Foundation through grant MCB-1449014, EPSCoR EPS-1004094, NSF CCF-1816969, and the National Institutes of Health (NIH) through grant 1-P20-GM113126-01. The material was based in part on work supported by the U.S. Department of Energy, Office of Science, under contract DE-AC02-06CH11357.

